# Structural modeling and thermostability of a serine protease inhibitor belonging to the Kunitz family from the tick Rhipicephalus microplus

**DOI:** 10.1101/2020.06.20.163246

**Authors:** Lívia de Moraes Bomediano Camillo, Graziele Cristina Ferreira, Adriana Feliciano Alves Duran, Flavia Ribeiro Santos da Silva, Wanius Garcia, Ana Lígia Scott, Sergio Daishi Sasaki

## Abstract

rBmTI-A is a recombinant serine protease inhibitor that belongs to the Kunitz-BPTI family and that was cloned from Rhipicephalus microplus tick. rBmTI-A has inhibitory activities on bovine trypsin, human plasma kallikrein, human neutrophil elastase and plasmin with dissociation constants in nM range. It is characterized by two inhibitory domains and each domain presents six cysteines that form three disulfide bonds, which contribute to the high stability of its structure. Previous studies suggest that serine protease inhibitor rBmTI-A has a protective potential against pulmonary emphysema in mice and anti-inflammatory potential, besides rBmTI-A presented a potent inhibitory activity against *in vitro* vessel formation. In this study, the tertiary structure of BmTI-A was modeled based on the structure of its *Sabellastarte magnifica* homologue. The structure stabilization was evaluated by molecular dynamics analysis. Circular dichroism data corroborated the secondary structure found by the homology modeling. Thermostability analysis confirmed the thermostability and the relation between the effects of the temperature in the inhibitor activity. The loss of activity observed was gradual, and, after 60 minutes of incubation at 90°C the inhibitor lost it completely.

## INTRODUCTION

Kunitz-BPTI-like inhibitors belong to a family of inhibitors that have one or more inhibitory domains characterized by a conserved spacing between cysteine residues, a typical disulfide binding pattern (Laskowski and Kato, 1980; Ranasinghe and McManus, 2013). Kunitz-BPTI-like domains are characterized by the conserved position of six cysteine residues that form three disulfide bonds contributing to the compaction and thermostability of these proteins (Brown et al., 1978; Ranasinghe and McManus, 2013). Different serine proteases can be inhibited by these molecules, such as trypsin, chymotrypsin, elastase, kallikreins, plasmin, factors XIa, IXa and cathepsin G (Ranasinghe and McManus, 2013).

The Kunitz domain has a secondary structural fold of the α/β type where the alpha helices and the β-sheet occur separately along the backbone. This domain is characterized by the presence of three highly conserved disulfide bridges that are necessary for stabilization of the native conformation of these inhibitors (Laskowski and Kato, 1980; Ranasinghe and McManus, 2013). The pancreatic bovine trypsin inhibitor (Bovine Pancreatic Trypsin Inhibitor - BPTI) is the most typical studied model of Kunitz-BPTI type inhibitors. BPTI has broad specificity and has inhibitory action towards serine proteases such as trypsin, chymotrypsin and elastase (Laskowski and Kato, 1980; Ranasinghe and McManus, 2013).

The standard mechanism of inhibition by a Kunitz type inhibitor involves a strong and non-covalent interaction like the complex that would be formed between the enzyme and its substrate. The Kunitz type inhibitors directly block the active site of the serine proteases. The segment responsible for protease inhibition is called the protease binding loop. This loop has a convex and prolonged structure, exposed to the solvent and is highly complementary to the enzyme concave active site. Residues that precede or follow this segment or from more remote regions can also participate in the interaction and influence the energy of association (Ascenzi et al., 2003).

Subsite nomenclature was adopted from Abramowitz et al. (1967) and Schechter et al. (1968) (Abramowitz et al., 1967; Schechter and Berger, 1968) work and used to describe enzyme specificities. According to their work, amino acid residues in a substrate undergoing cleavage are designated P1, P2, P3, P4 and so on in the N-terminal direction from the cleaved bond. The P1 position is the main determinant of the specificity of protease recognition by canonical inhibitors. A large portion of the protease-inhibitor contact is made only by the residue from the P1 position, which can penetrate deeply into the reactive protease cavity. Kunitz-BPTI type inhibitors for trypsin typically have the Arg and Lys residues at their P1 sites (Grzesiak et al., 2000; Ranasinghe and McManus, 2013) while Kunitz type inhibitors for chymotrypsin have Leu and Met at their P1 sites (Grzesiak et al., 2000).

Kunitz-BPTI like serine protease inhibitors present in Rhipicephalus microplus tick are known as BmTIs. They have one or more Kunitz-BPTI domains. These inhibitors were initially purified from larvae and egg of ticks(Andreotti et al., 2002; Sasaki et al., 2004). The BmTI-A inhibitor was initially obtained from the extraction of Rhipicephalus microplus tick larvae (Tanaka et al., 1999). The BmTI-A cDNA was cloned from the gut of ingurgitated females and inserted into a pPIC9K expression vector, its expression was carried out in Pichia pastoris system and the purification was performed by trypsin-sepharose chromatography (Soares et al., 2016). The recombinant rBmTI-A obtained has two Kunitz-BPTI domains, with a total molecular mass of 13.87 kDa. Domain 1 has the amino acid Arg at P1 site with an inhibitory action against trypsin and human plasma kallikrein (HuPK). Domain 2 has the amino acid Leu at P1 site with an inhibitory action against human neutrophil elastase (HNE) (Sasaki et al., 2004; Soares et al., 2016). The amino acid residues sequence of the recombinant inhibitor domains is shown in Fig. 1.

**Fig. 1.**
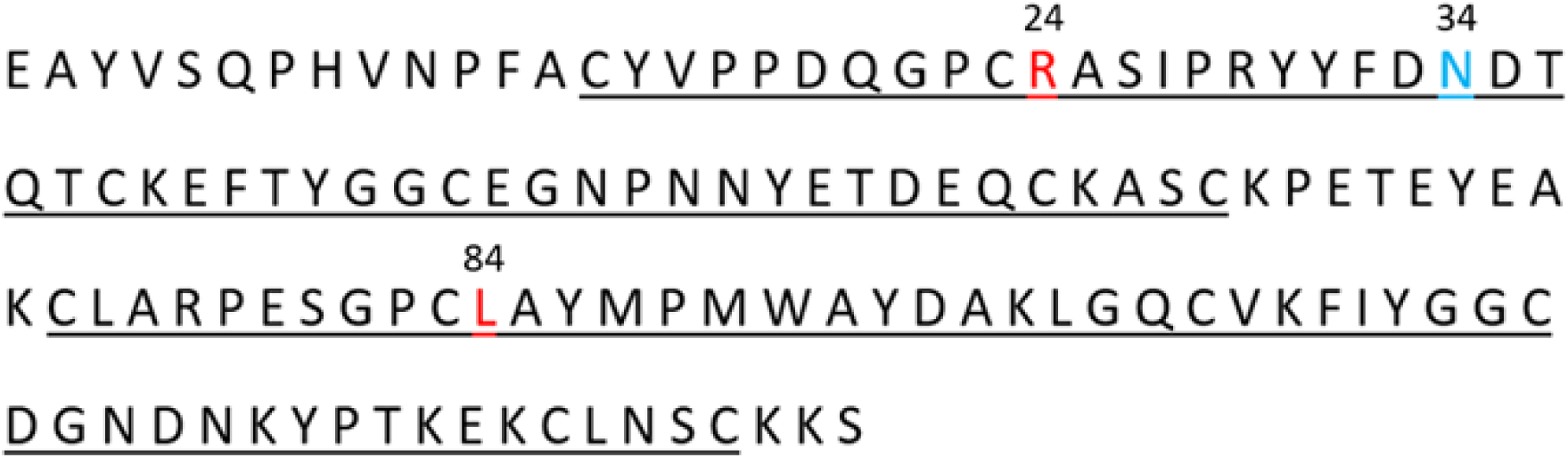
Sequence of rBmTI-A inhibitor. The regions of each Kunitz-BPTI domain are delimited by the underlined amino acids. Amino acids from P1 site from each domain are marked in red: amino acid arginine at position 24 and amino acid leucine at position 84. Amino acid at position 34 indicates the potential asparagine glycosylated, highlighted in blue. Theoretical molecular mass of the inhibitor: 14.28 kDa. Total number of residues: 127 (Lourenço et al., 2018, 2014; Soares et al., 2016).

rBmTI-A presents inhibitory activities on bovine trypsin, human plasma kallikrein and human neutrophil elastase (Sasaki et al., 2004; Soares et al., 2016; Tanaka et al., 1999). Serine proteases are involved in several important physiological processes, such as protein metabolism, digestion, blood coagulation, apoptosis, regulation of the development of immunity, inflammatory processes and fertilization (Di Cera, 2009; Hedstrom, 2002; Pham, 2006). Serine proteases are classified according to the specificity of their substrate and the most common types are trypsin-like, subtilisin-like, chymotrypsin-like, elastase-like, kallikreins and cathepsins (Di Cera, 2009; Page and Di Cera, 2008). Since serine proteases are involved in several physiological and pathological processes, the study of their inhibition is very import as a promise of therapeutic use for the treatment of diseases such as cancer, neurodegeneration, tissue inflammation and infections. At least 23 different families of canonical inhibitors have been identified based on structural criteria (Patston, 2000) and two of the most studied families are serpins and Kunitz-BPTI inhibitors.

Previous studies suggest that serine protease inhibitor rBmTI-A has a protective potential against pulmonary emphysema in mice, attenuating the characteristic effects of disease onset and progression (Lourenço et al., 2018, 2014) In addition, rBmTI-A is effective in attenuating the changes in lung mechanics, inflammation and remodeling of the airways in an animal model of chronic allergic pulmonary inflammation (Florencio et al., 2015). Soares et al. (Soares et al., 2016) demonstrated a possible role of rBmTI-A inhibitor in controlling angiogenesis in human umbilical vein endothelial cell (HUVEC) line. In this study, rBmTIA was able to disrupt vessel formation by inhibiting plasma kallikrein, neutrophil elastase and plasmin. The inhibition of plasma kallikrein led to the blocking of bradykinin release, which is responsible to stimulate angiogenesis and vascular modeling. The same study has also shown that the inhibition of plasma kallikrein, neutrophil elastase and plasmin led to the blocking of other growth factors release, like VEGF (vascular endothelial growth factor), FbGF (fibroblast growth factor) and TGF-b1 (transforming growth factor), inhibiting vessel formation and blocking wound healing which is favorable to the tick feeding.

Considering the importance of the action of rBmTI-A inhibitor against emphysema and its anti-inflammatory and anti-angiogenic potentials, it is equally important to understand its structure and how it can interact with molecules of interest in a variety of diseases. Until now, there is no report about its three-dimensional structure in literature. This study presents a reliable homology model for rBmTI-A structure, corroborated by circular dichroism spectroscopy analysis. Thermostability assays were performed to test inhibition activity and structural stability in different temperatures.

## MATERIAL AND METHODS

### rBmTI-A structure modeling

Basic information about the serine protease inhibitor was obtained from previous study using the rBmTI-A amino acid sequence (Soares et al., 2016; Tanaka et al., 1999). Crystallographic model of *Sabellastarte magnifica* hydrolase inhibitor SmCI (4BD9, chain B) was used as template to the theoretical rBmTI-A model, according to data obtained from HHpred (Soding et al., 2005), available at Max-Planck Institute for Development Biology server (https://toolkit.tuebingen.mpg.de/tools/hhpred). The initial theoretical model was generated by MODELLER v9.23 using homology modeling method (Baker, 2001; Cavasotto and Phatak, 2009; Ginalski, 2006; Xiang, 2006).

### Molecular dynamics simulation

Theoretical model obtained by MODELLER was submitted to molecular dynamics (MD) simulation in GROMACS (Groningen Machine for Chemical Simulation) v5.1 (Van Der Spoel et al., 2005) in the presence of explicit water molecules. All MD simulations were performed with a rectangular 10 Å solvent box under constant temperature (298,15 K) and pressure (1 atm). Protonation states of charged groups at pH 7.0 were obtained by PROPKA v3.0 of PDB2PQR v2.0 server (Dolinsky et al., 2004), available at http://server.poissonboltzmann.org/pdb2pqr_2.0.0. The charges of the model were neutralized by the addition of Na^+^ and Cl^-^ ions to maintain neutrality by ion placing method Monte Carlo. Force field AMBER99SB (Showalter and Bru, 2007) and TIP3 water model were used. Energy minimization to generate the starting configuration of the system was performed using steepest descent method (Troyer and Cohen, 1995). A first MD step of 200 ps with position restrains was applied to relax the system and then 50 ns of unrestrained MD simulation were applied to evaluate the stability of the structure.

### Evaluation of theoretical rBmTI-A model

The overall stereochemical quality of the model was tested with MolProbity v4.5.1 (Williams et al., 2018), ProSA-web (Wiederstein and Sippl, 2007), available at https://prosa.services.came.sbg.ac.at/prosa.php and Verify3D (Eisenberg et al., 1997), available at https://servicesn.mbi.ucla.edu/. Model secondary structure content was obtained with Quick2D (Biegert et al., 2006) and STRIDE (Heinig and Frishman, 2004).

### Preparation of the rBmTI-A

The recombinant inhibitor (rBmTI-A) was obtained using the same protocol described in the previous study (Soares et al., 2016). Briefly, the rBmTI-A cDNA was cloned into a pPIC9K plasmid and it was inserted into Pichia pastoris yeast. The protein expression was performed during five days with 0.5% of methanol to induce it. After this period the supernatant was centrifuged 1,500 rpm x 30 min. The purification was performed using the trypsin-Sepharose column. The purified samples were desalted using 50 mL Amicon system, 3 kDa cut-off and samples were stored in phosphate saline buffer, pH 7.4.

### Circular dichroism spectroscopy

An aliquot of rBmTI-A was prepared for circular dichroism (CD) spectroscopy and thermostability experiments. The concentration used was 0.36 mg/mL in sterile phosphate saline buffer, pH 7.4. Phosphate saline buffer was used as control. For simple verification, the inhibitory activity was tested. CD spectra were measured using a Peltier coupled Jasco J-815 spectropolarimeter for temperature control. Thermal stability was monitored at temperatures from 20 °C to 90 °C at 10 °C intervals, using a 0.1 cm quartz cuvette. The spectra were obtained in wavelength ranging from 195 to 260 nm. The reads were performed after 5 min in each interval. After it reached the temperature of 90 °C, a new read was made at low temperatures (40 °C and 20 °C) to verify if the structure would return to its initial conformation. K2D3 software (Louis-Jeune et al., 2012) was used to estimate the secondary structure of the protein by the CD spectrum.

### Thermostability assays

The rBmTI-A concentration used was 5.8 nM in PBS (Phosphate buffer solution). Aliquots were made in triplicate and incubated in a thermal cycler (T100 thermal cycler Bio Rad) at different times and temperatures. After completion of the 30 min or 60 min incubation, the inhibitor was immediately incubated on ice until enzyme activity tests. Inhibitory activity was monitored using bovine trypsin and fluorogenic substrate Z-Phe-Arg-MCA, in 100 mM Tris-HCl buffer, pH 8.0 containing 0.15 M NaCl at 37 °C. Enzyme and inhibitor were incubated for 10 min at 37 °C before addition of the substrate. The final volume of the assay was 100 μl. The monitoring of the results was performed by relative fluorescence units (RFU) readings using Synergy HT Multi-Mode Microplate Reader Biotek®. The following parameters were used: gain 65; optic exposition Top; wavelength 320/20 (excitation) and 460/40 (emission). The residual enzyme activity was calculated from the ratio between the RFU obtained of assays containing the inhibitor, enzyme and substrate and RFU obtained of assays containing only enzyme and substrate. We obtained the inhibitory activity value (IA) subtracting the value of residual enzyme activity (RA) from 1 (IA: 1-RA). The analysis of these results was performed using GraphPad Prism 5.01 software (Swift, 1997).

## RESULTS

### rBmTI-A model

The final three-dimensional model obtained for rBmTI-A inhibitor is shown in Fig. 2. Fig. 2A shows the model colored by secondary structure where α-helix are represented by red, β-strands by yellow and coiled coils by green. In Fig. 2B, the two functional domains are colored in gray. Amino acids of P1 site of both domains are shown as yellow sticks, Arg at position 24 of the first domain and Leu at position 84 of the second domain. Cysteine residues forming disulfide bonds are represented by lemon green sticks. The average percentage for secondary structure of rBmTI-A model is 17% α-helix, 23% β-strands and 60% coiled coils. The structure model presented a good overall stereochemical quality with 92.7% of its residues in most favored regions and 99.1% in allowed regions of the Ramachandran plot (Kleywegt and Jones, 1996; Zhou et al., 2011). All residues presented negative potential energies and the Z-score of the model is -5.32. Verify 3D test resulted in 85.79% of the residues with an averaged 3D/1D score >= 0.2. The final structural rBmTI-A model presented a RMSD (Root Mean Square Deviation) stabilization after 1000 ps of MD simulation, with positional deviation of atoms of 0.242909 nm. These findings indicate that the final theoretical model of rBmTI-A structure after MD simulation presents high stability and it is reasonable for further studies.

**Fig. 2.**
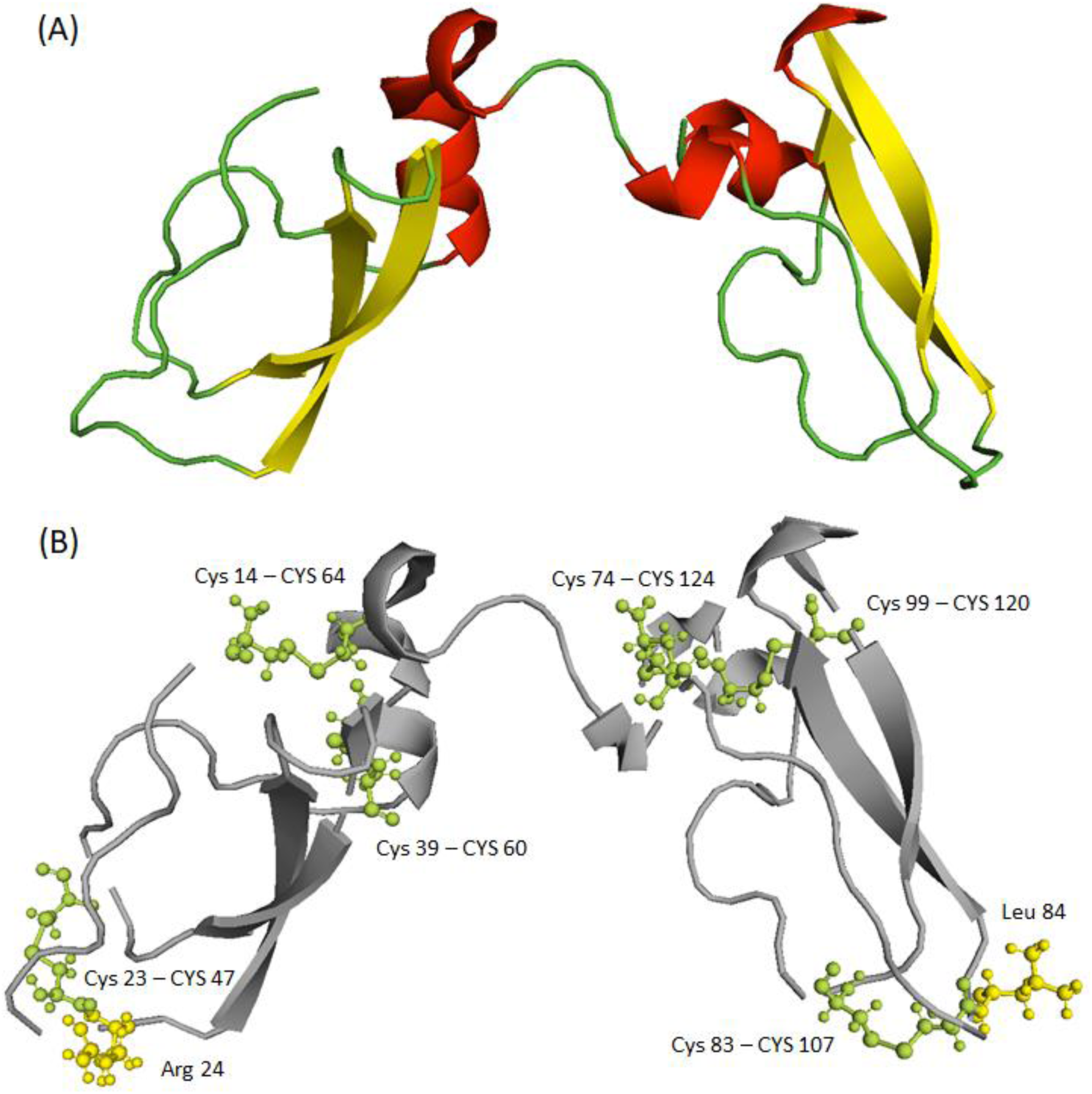
Three-dimensional structure of rBmTI-A inhibitor. A) rBmTI-A model colored by secondary structure: α-helix are represented by red, β-strands by yellow and coiled coils by green. B) The two functional domains are colored in gray. Amino acids of P1 site of both domains are shown as yellow sticks, Arg at position 24 of the first domain and Leu at position 84 of the second domain. Cysteine residues forming disulfide bonds are represented by lemon green sticks.

### Circular dichroism (CD) spectroscopy and thermostability assays

The Fig. 3 shows the rBmTI-A CD spectrum. Between 20 °C and 60 °C it was not observed any significant change in the inhibitor secondary structure and the spectrum of this temperature range is represented as one curve at 20 °C. The CD spectrum at 20 °C is characteristic of proteins with high irregular secondary structure content, as indicated by the measure around 200 nm, which is typical of coil regions. At 70 °C the first modifications in the inhibitor structure are noticed and at 80 °C to 90 °C the most significant changes are observed, indicating loss of the original structure of the inhibitor. Temperature reduction after denaturation does not recover the original spectrum, indicating that the process was irreversible after heating the structure at 90 °C. CD analysis of the secondary structure content at 20 °C using K2D3 deconvolution spectrum software version 1.0 (Louis-Jeune et al., 2012) showed that it is composed by 13% α-helix, 28% β-strands and 59% coiled coils.

**Fig. 3.**
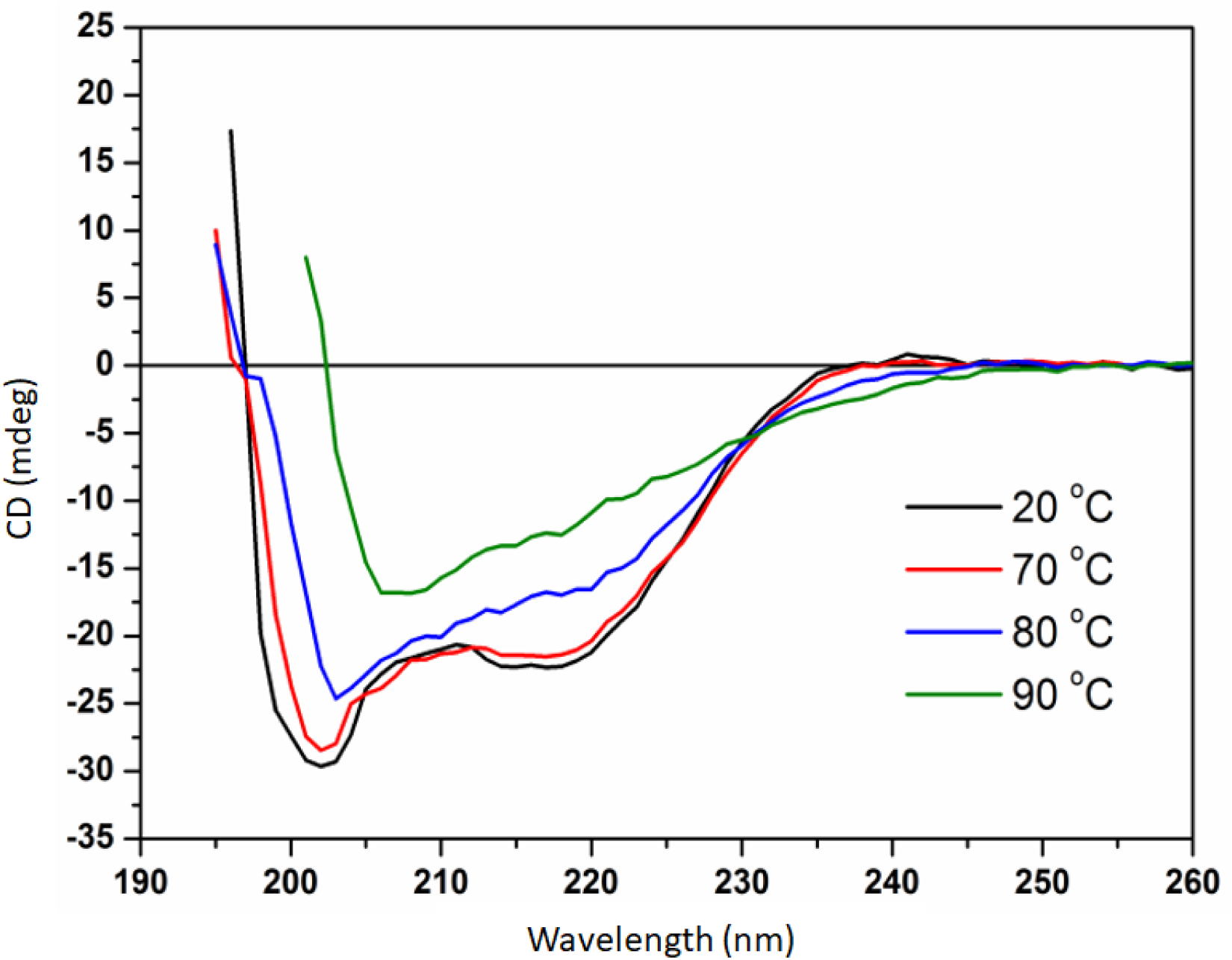
CD spectrum of rBmTI-A structure. Between 20°C and 60°C it was not observed any significant change in the inhibitor structure and the spectrum of this temperature range is represented as one curve at 20°C. At 80°C and 90°C, the structure loses its stability, therefore losing its functional state.

The results in Fig. 4 show the inhibitory activity of trypsin enzyme after readings with the inhibitor incubated at different temperatures and times (Fig. 4A incubation time of 30 min and Fig. 4B incubation time of 60 min). The loss of activity observed was gradual compared with the sample incubated at 4 °C and at 90 °C the inhibitor lost its activity completely for both incubated times. At 70 °C, incubated for 30 min, the reduction in inhibitory activity was 42% compared to 54% reduction when the incubation time at the same temperature was 60 min. It suggests that loss of activity of rBmTI-A inhibitor is related to not only the temperature, but also to the incubation time.

**Fig. 4.**
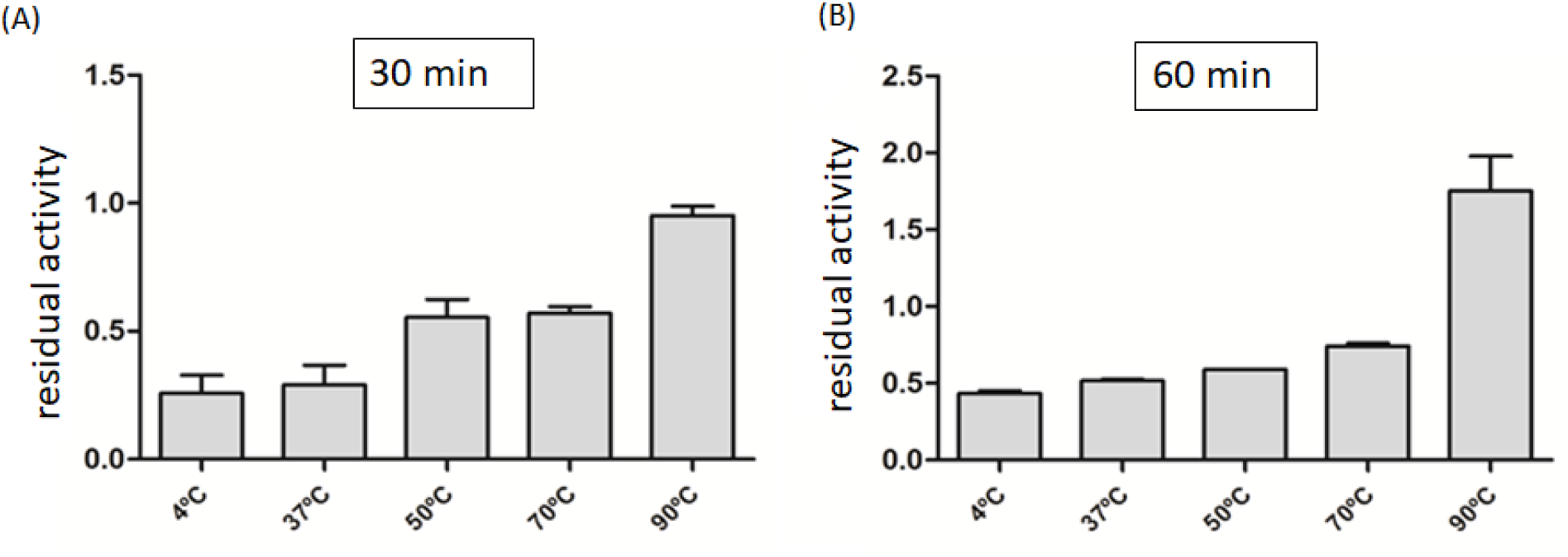
Molecule structure stability at different temperatures. The rBmTI-A inhibitor (5.8 nM) was incubated for A) 30 minutes and B) 60 minutes, before the enzymatic tests. Substrate: Z-Phe-Arg-MCA final concentrate 0.06mM. Enzyme: 0.25ng/μl. The final volume the assay was 100 μl **/*** ANOVA analysis followed by Turkey test comparing different temperatures. The statistical difference with p < 0.05 of the reduction of the inhibitory activity compared to 4°C.

## DISCUSSION

rBmTI-A is a serine protease inhibitor belonging to the Kunitz I2 family (Kunitz and Northrop, 1936; Rawlings et al., 2004) from Rhipicephalus microplus with inhibitory activities on bovine trypsin, HuPK, HNE and plasmin with dissociation constants (Ki) in nM range. It is characterized by two inhibitory domains and the presence of six cysteines residues that form three disulfide bonds in each domain, which contribute to the high stability of its structure (Sasaki et al., 2004; Soares et al., 2016). Several protease inhibitors have been characterized from Rhipicephalus ticks. Most of these inhibitors belong to the Kunitz-BPTI family and come from different tissues and stages of organism development, such as RmKK (R. microplus tissue kallikrein), purified from R. microplus eggs with inhibitory activity on tissue kallikrein (Abreu et al., 2014), BmCI (B. microplus chymotrypsin inhibitor) from fat body and hemocyte cDNA libraries of R.micropulus with inhibitory activity towards bovine chymotrypsin (Lima et al., 2010), RsTI (R. sanguineus trypsin inhibitor) from R. sanguineus larvae that inhibits serine proteases such as bovine trypsin, human neutrophil elastase and HuPK (Azzolini et al., 2003) and others Rhipicephalus microplus subtilisin inhibitors (BmSI), BmSI-7 and BmSI-6 purified from eggs, with activity on neutrophil elastase and subtilisin A (Sasaki et al., 2008).

The Ramachandran plot is a diagram of φ versus ψ backbone dihedral angles for each residue in the protein. The diagram is divided into favored, allowed, and disallowed regions and it is a simple way to access the quality of a protein structure without any experimental data (Kleywegt and Jones, 1996; Zhou et al., 2011). Regions of the plot are defined for general case and for especial case of the amino acid isoleucine and valine, glycine, pre-proline, trans proline and cis proline. According to the literature (Kleywegt and Jones, 1996; Zhou et al., 2011), for a high-quality model it is expected ∼ 80% of the residues to be in most favored regions of the plot. The Ramachandran plot for the rBmTI-A model has 92.7% of its residues in most favored regions.

ProSA is a quality check that discriminate correct fold and incorrect fold in a three-dimensional protein model based on statistical analysis of all available protein structures in PDB and energy plots as a function of amino acid sequence. In both analysis, negative values indicate a high-quality model (Wiederstein and Sippl, 2007). Verify 3D is capable to determine the compatibility of a three-dimensional protein model with its own amino acid sequence to test model accuracy (Eisenberg et al., 1997). ProSA and Verify 3D analysis show that the structure obtained for rBmTI-A is a high-quality model. The predicted secondary structure for the model was corroborated by the circular dichroism analysis, indicating the presence of high content of coil regions and β-strands secondary structure. These coiled coil regions are important because they are strictly related to the correct assembly of the protein final and functional structure (Linding et al., 2003; Phillips, 1992). Some studies suggest that these regions have been known as linkers, probes and inhibitors of protein-protein interactions (Adamson et al., 1993; Watkins et al., 2015).

The trajectory of a molecular dynamic simulation is the representation of structural dynamics of the proteins along the time and it is useful for sampling the conformational space and study local and global motions of macromolecules. The RMSD value in this context is a standard measure of the structural distance between the coordinates, an average distance between groups of atoms. The RMSD between two points in the trajectory time is a measure of how much the conformation of the protein has changed (Brüschweiler, 2003). For the structure of the rBmTI-A model, the fluctuation of the values of total RMSD are around 0.24 nm indicating that the structure of the inhibitor model deviated steadily from its initial structure, towards a more energetically stabilized one.

Kunitz inhibitors can be single domain or multiple domains. Bikunin and Boophilin are examples of serine protease inhibitors with Kunitz domain repeated twice (Delaria et al., 1997; Macedo-Ribeiro et al., 2008). Penthalaris and Ixolaris are tissue factor pathway inhibitors (TFPIs) obtained from the Lyme disease vector tick saliva, Ixodes scapularis. Both inhibitors have multiples Kunitz-type domains repeated, twice in Penthalaris and five times in Ixolaris (Francischetti et al., 2004, 2002). Other inhibitors can have up to 12 repeated domains, as the proteinase inhibitor from the hookworm Ancylostoma caninum (Hawdon et al., 2003). The repeated domains may be able to interact independently with multiple proteases at their reactive sites at the same time. Boophilin is a good example of a multivalent Kunitz-like inhibitor, it can interact with serine proteases in a canonical manner and with thrombin in a non-canonical way at the same time (Macedo-Ribeiro et al., 2008). Ixolaris first domain, with amino acid Glu at the P1 site, has inhibitory activity towards FVIIa (blood-coagulation factor VIIa) catalytic site, while second domain is responsible for the inhibitory activity towards FXa (blood-coagulation related factor Xa) (Francischetti et al., 2002). However, studies with Penthalaris showed that not always all the separated domains will interact at the same time or contribute for the inhibitory process. Domains 1, 2 and 5 have, respectively, Arg 26, Arg 87 and Lys 253 at their P1 site and appear to interact with FVIIa (blood-coagulation factor VIIa) catalytic site. Domains 3 and 4 have, respectively, amino acids Gly 145 and Asp 199 at their P1 site and do not show ideal catalytic recognition or contribution for the inhibitory activity to occur (Francischetti et al., 2004). In the context of pulmonary emphysema, rBmTI-A could act as a multivalent inhibitor, since elastase e other proteases that can interact with its domains, such as kallikreins, are involved in the inflammatory process (Shaw and Diamandis, 2007).

The presence of the characteristic Kunitz-type conserved spaced cysteines forming structural stabilization disulfide bonds is very important for the inhibitory process to occur (Bode and Hurder, 1992; Ranasinghe and McManus, 2013). The disulfide bonds are required for the maintenance of the functional structure of the inhibitor. Classical C1–C6 and C3–C5 disulfide bonds are related to the conservation of the structure final conformational and the C2–C4 bond is responsible to stabilize the domains (Laskowski and Kato, 1980; Ranasinghe and McManus, 2013). The Kunitz-type domains found in Penthalaris and Ixolaris present loss of some of these disulfide bonds. Penthalaris domains 1, 2 and 5 and Ixolaris second domain have only two out of three of the expected conserved disulfide bonds. The absence of these bonds produces a more flexible loop structure at P1 sites, leading to difficulties in the interaction with the target protease catalytic site. But it does not mean that these domains cannot interact at all. They have inhibitory activity, as it was cited before, but this activity often occurs in other sites than the catalytic ones (Demchenko, 2001; Francischetti et al., 2004; Ranasinghe and McManus, 2013).

Several factors can modify the structure of a protein, leading to denaturation and activity loss, such as pH and temperature variations (Davis and Williams, 1998; Dill and Shortle, 1991; Privalov, 1990; Tanford, 1968; Yang and Honig, 1993). One of the techniques used to analyze this structural change during protein denaturation is circular dichroism spectroscopy (Greenfield, 2006; Johnson, 1990; Kelly and Price, 2000; Woody, 1995). In some Kunitz-type inhibitors, the denaturation caused by temperature increasing can be reversible depending on the temperature range. Soy trypsin inhibitor incubated at 70 °C loses its functional structure. When it is incubated at 70 °C and then back to 30 °C, it is capable to recover its activity. But the process became irreversible when it is incubated at 90 °C (Kunitz and Northrop, 1936), as the data of CD and thermostability assays also confirmed to rBmTI-A inhibitor. Inhibitors of other ticks have also been characterized by their thermostability. The Kunitz-type HlChI (Haemaphysalis longicornis chymotrypsin inhibitor) has lost its activity at pH 10 and 70 °C, but when the inhibitor was incubated at pH 4 and 100 °C, it could maintain its activity. Studies with thermostable plant Kunitz-type inhibitors show that activity loss can also result from the addition of reducing agents, such as DTT (dithiothreitol), by losing or reducing the disulfide bonds stability and inhibitory activity (Brandão-Costa et al., 2018; Chan et al., 2014; Fang et al., 2010; Konigsberg, 1972). The structure obtained for the rBmTI-A inhibitor with the cysteine residues forming the disulfide bonds conserved, corroborates the thermostability evaluated by the circular dichroism experiment and by the inhibitory activity assay. The rBmTI-A inhibitor is a stable inhibitor at high temperatures, allowing the inhibitory activity to be observed up to at least 70 °C.

## CONCLUSIONS

This study presented a reliable homology model of the structure of BmTI-A inhibitor, corroborated by the circular dichroism spectroscopy analysis. This structure is extremely important for its inhibitory activity. As shown by circular dichroism data, any modification on the structure of the inhibitor caused by the increase of the temperature leads to a reduction of its inhibitory activity. We suggest that the thermostability of rBmTI-A is related to the presence of cysteines residues forming three disulfide bonds in each domain of its structure.

## ACKNOWLEDGMENTS

We are grateful to Fundação de Amparo à Pesquisa do Estado de São Paulo (FAPESP); Conselho Nacional de Desenvolvimento Científico e Tecnológico (CNPq) – Brazil; and Coordenação de Aperfeiçoamento de Pessoal de Nível Superior (CAPES) for supported this work.

