## supplemental figures for "Structural modeling and thermostability of a serine protease inhibitor belonging to the Kunitz family from the tick Rhipicephalus microplus"

### SUPPLEMENTARY MATERIALS

#### rBmTI-A model quality results

The structural model was submitted to quality tests. MolProbity provides quality validation of 3D structures of proteins through detailed all-atom contact analysis (Chen et al., 2010). It calculates Ramachandran plots and geometry statistics of the structure. Fig. 1 shows the Ramachandran plots for rBmTI-A. 92.7% of the residues are in most favored regions and 99.1% in allowed regions, one outlier was found for the general case. The amino acid glutamine of the position 69 is in a disallowed region. Cβ deviation is a measure of problematic geometry that often indicates incompatibility between side chain and main chain conformations. Deviations of ≥ 0.25Å are correlated with local inappropriate fitting of the model (Lovell et al., 2003). The model presented one amino acid with Cβ deviations ≥ 0.25Å. MolProbity also analyses bad bonds and bad angles in the structure. It is expected that a model or experimentally determined structure should have < 0.1% of bond angles that differ from the accepted values (Engh and Huber, 2012). No bad bonds were observed, but 10/1221 (0.82%) bad angles were found for rBmTI-A model. The Fig. 2A and Fig. 2B shows the results for rBmTI-A model ProSA evaluation and Fig 3 shows the Verify3D test. It resulted in 85.79% of the residues with an averaged 3D/1D score >= 0.2.

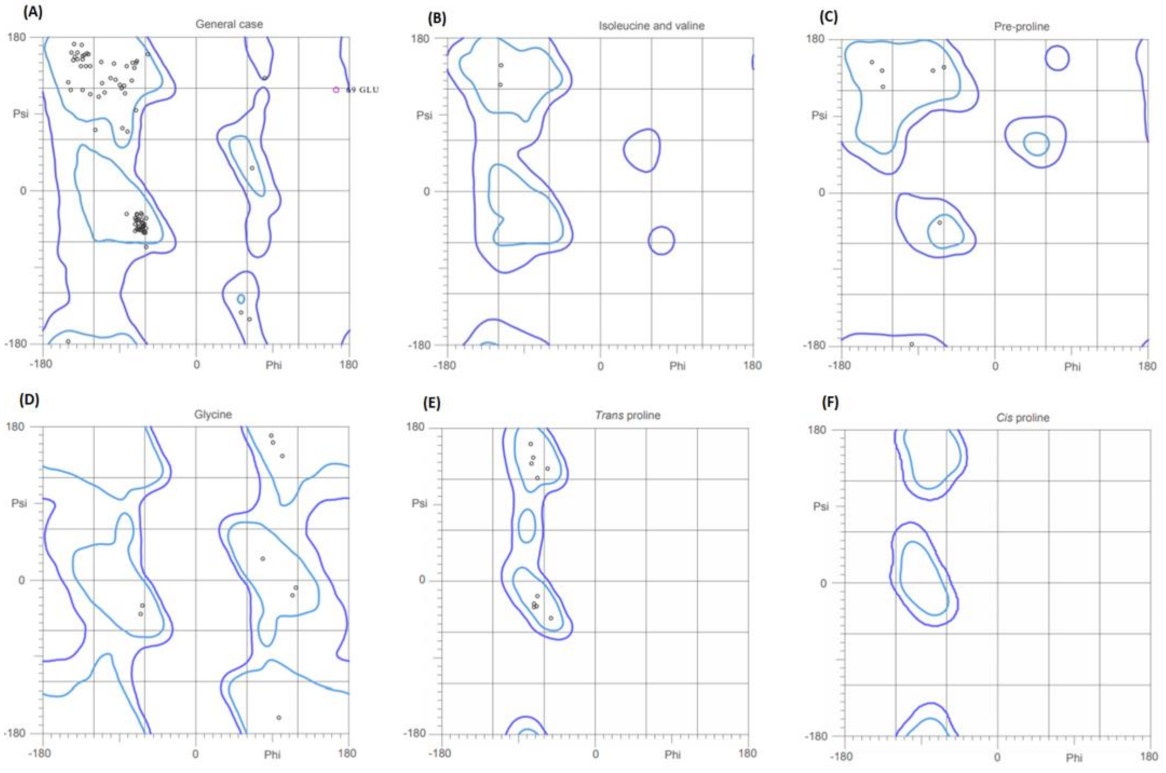


**Fig. 1**. **Ramachandran plots for rBmTI-A model.** A) General case. B) Isoleucine and valine. C) Pre-proline. D) Glycine. E) Trans proline. F) Cis proline. 92.7% of the residues are in most favored regions and one outlier were found in the general case plot (glutamine).

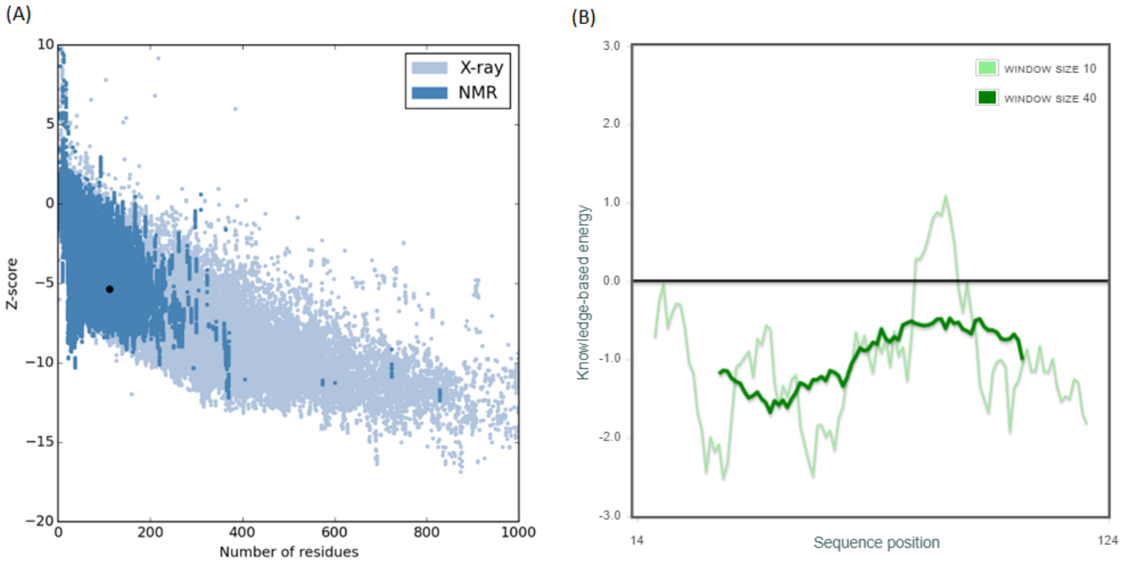


**Fig. 2. ProSA quality check.** A) ProSA Z-scores for rBmTI-A (black dot). Light blue and blue dots are Z-scores of PDB structures determined by X-ray crystallography and NMR. B) Energy plot for rBmTI-A. Negative values indicate a high-quality model.

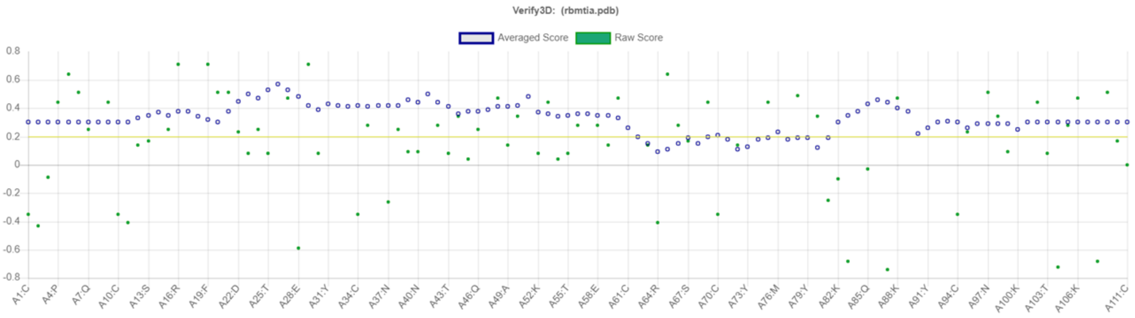


**Fig. 3. Verify3D score profile calculated for rBmTI-A model.** At least 80% of the scores >= 0.2 indicate a high-quality model. rBmTI-A model presented 85.79% of the score >= 0.2.
